# A comprehensive and comparative study on the action of pentacyclic triterpenoids on *Vibrio cholerae* biofilms

**DOI:** 10.1101/2020.01.06.896183

**Authors:** Sudipta Paul Bhattacharya, Arijit Bhattacharya, Aparna Sen

## Abstract

While serving as environmental reservoir for *V. cholerae* infection, biofilms are also crucial for intestinal colonization of the pathogen. Triterpenoids, a group of bioactive phytochemicals, have been tested for antibiofilm activity against model biofilm forming bacteria in recent times. In this context, glycyrrhetinic acid (GRA), ursolic acid (UA) and betulinic acid (BA), representing three categorically distinct groups of pentacyclic triterpenoids, are targeted for profiling their impact on *Vibrio cholerae* C6709 biofilms. The triterpenoids substantially affected biofilm associated attributes like formation, substratum adherence and dispersion from preformed biofilms. Though at variable degree, the compounds decreased cell surface hydrophobicity and composition in terms of macromolecular content. Not only EPS-associated extracellular enzyme activities were estimated to be reduced by triterpenoid exposure, ultra structural analysis also revealed that GRA, UA and BA can affect extracellular polymeric substance (EPS) content. Albeit total extracellular proteolytic activity remained unaffected by the triterpenoids, GRA treatment resulted in considerable reduction of extracellular gelatinase activity. Molecular docking analysis indicated potential interaction with cyclic di-GMP sensor VpsT, autoinducer-2 sensor kinase LuxP-LuxQ and transcriptional activator HapR, component of complex quorum sensing networks modulating biofilm formation. Comprehensive analysis of antibiotic action revealed accentuation of cephalosporin antibiotics with GRA and UA while BA potentiated action of fluoroquinolones, widening the scope of combinatorial therapeutic strategy.

## Introduction

Biofilm has been defined as an assemblage of microbial cells surrounded by a matrix of extracellular polymeric substances (EPS) secreted by those cells. Biofilm formation can serve as a survival mechanism for bacteria as it can provide protection from toxic compounds, such as antibiotics and microbicides (Hall-Stoodley and Stoodley, 2009) beside protecting from various environmental challenges such as pH, salinity, and metal toxicity (Koo et al., 2017).

The EPS matrix provides a physical structure that segregates in microdomains (Lawrence et al., 2007) and it’s composition is critical for biofilm integrity which in turn plays pivotal role in biofilm development, dispersion and susceptibility to antimicrobials (Flemming and Wingender, 2010). As per reports from Centres for Disease Control and Prevention (CDC) and National Institutes of Health (NIH), the estimated frequency of infections caused by biofilms lies between 65% and 80% respectively (Moser et al., 2018).

Even with significant progress in the management of diarrhoeal diseases like cholera has been achieved with improved hygiene, development of new antimicrobials and vaccines, the burden remains the same, especially in neonates and infants (http://www.who.int/cholera/en). In case of cholera, though oral rehydration treatment remains the mainstay, antimicrobial therapy is implemented at times to curb the duration of the ailment which is complicated with emergence of *V. cholerae* strains resistant to major antibiotics (Ghosh and Ramamurthy, 2011).

*V. cholerae* biofilm in aquatic environments serve as a potential reservoir of pathogenic cells causing periodic occurrences of the severe diarrhoeal disease, typically resulting from consumption of pathogen contaminated drinking water. One key factor for environmental transmission is biofilm formation providing a substantial survival advantage to aquatic organisms (Mourino-Perez et al., 2003). Deciphering the molecular mechanism for biofilm development identified VpsR and cyclic-di-GMP mediated pathways as the key modulators in *V. cholerae* (Hsieh et al., 2018). Biofilm formation by the pathogen is directly linked to its pathogenesis (Silva and Benitez, 2016) and increased antibiotic resistance has been shown in biofilms of *V. cholerae* O139 cells (Gupta et al., 2018). Sultana et al. (2018) recently showed biofilms to be a mean of persistence and an integral component of the annual life cycle of toxigenic *V. cholerae* as well (Sultana et al., 2018).

Strategies to develop antibiofilm agents against diverse groups of microorganisms including *V. cholerae* include designing of diguanylate cyclase targeting molecules (Sambanthamoorthy et al., 2012), development of quorum sensing antagonists targeting multiple effectors in *V. cholerae* (Faloon et al., 2010; Kim et al., 2018) etc. Pentacyclic triterpenoids have been implemented in antibiofilm research recently (Gilabert et al., 2015; Kannan et al., 2019; Rajkumari et al., 2018). In fact, 20 different compounds with triterpenoid scaffold have been screened for antibiofilm activity against Gram positive pathogens (Silva et al., 2019). Pentacyclic triterpenoids are classified into three classes, namely, lupane, oleanane or ursane (e Silva Mde et al., 2012). In this study, with a target to generate comparative profile, representative molecules from each of the group namely glycyrhetinic acid (GRA), ursolic acid (UA) and betulinic acid (BA) are tested for their impact on biofilm formation, dispersion and motility. Biofilm integrity and composition analysis indicated significant alteration in EPS production and composition. Possible interaction between the triterpenoids and a series of antibiotics was also evaluated against planktonic cells and specific antibiotic-triterpenoid combinations were tested against *V. cholerae* C6709 biofilms. Possible influence on quorum sensing was also indicated by molecular docking between the triterpenoids and major quorum sensing regulators like VpsT, LuxP, LuxQ and HapR.

## Materials and methods

### Selection of representative triterpenoid, bacterial strains and growth condition

Representative triterpenoids from each of the three major groups, namely betulinic acid (BA) from lupine; glycyrrhetinic acid (GA) from β-amyrin and ursolic acid (UA) (Sigma) from ursane families of triterpenoids were selected. For the studies batch cultures of *Vibrio cholarae* C6709 strains (bacterium O1, biotype El tor, serotype Inaba) were grown at 37°C with shaking in Luria-Bertani (LB) or Mueller-Hinton medium (for antibiotic assays). Swarming motility assay was carried out in LB medium supplemented with 0.5% agar, as described elsewhere (Rashid et al., 2000).

### Determination of Minimum Bactericidal Concentration

Minimum bactericidal concentration (MBC) assay was performed essentially as described by Provenzano et al. with some modifications (Provenzano et al., 2000). The MBC was defined as that at which no viable bacteria was recovered. For each phytochemical-treated system respective solvent-control was used.

### Biofilm Formation Assay

Biofilms were allowed to form into borosilicate glass test-tubes after an incubation of 24 h at 37°C. The tubes were then rinsed with distilled water and filled with crystal violet stain. After 5 min, the tubes were rinsed. The biofilm-associated crystal violet was suspended with ethanol (70%), and the optical density (OD) of the resulting suspension was measured.

### Biofilm-Dispersion assay

Biofilms were grown for 24 hrs in the LB medium. Aged medium was replaced with fresh medium containing phytochemicals to induce biofilm dispersion that was measured by an increase in turbidity for media at 600 nm.

### Microscopic examination of integrity of Biofilms

The biofilm integrity was examined for detecting minute alterations in matrix structure by scanning electron microscopy (SEM).

### Autoaggregation assay

Bacteria were grown at 37°C for 24h in LB broth. The cells were harvested by centrifugation and suspended in PBS to 0·5 OD units at 600nm. 2 ml bacterial suspension was placed in each tube, centrifuged and then resuspended in PBS along with the triterpenoids. After 2-3h exposure at 37°C, 1 ml of the upper suspension was transferred to another tube and the OD was measured. Aggregation was expressed as 1−(OD_upper suspension_/OD_total bacterial suspension_)×100.

### EPS extraction and assay

To investigate EPS composition and properties, *V. cholerae* strains were cultivated as agar-grown biofilms at 37°C for 24 h. After carefully scraping the bacterial biomass from the agar surface and suspending in NaCl, the cells were dispersed by vigorous stirring followed by high-speed centrifugation to separate it from the EPS. The supernatant was filtered through cellulose acetate membranes to remove residual bacteria and yield a cell-free EPS solution.

### Chemical composition analysis

The major structural groups of the partially purified EPS were detected in absorption mode with Fourier-transformed infrared (FT-IR) spectroscopy using FTIR spectrometer Perkin-Elmer Spectrum 100 and the spectra were obtained in the wave number range of 1000-3500 cm−1 using Spectrum software. Total protein content was measured by adding trichloroacetic acid (TCA) (final concentration, 12%) to the EPS solution, incubating the mixture on ice for 30 min before centrifugation at 13000 rpm for 20 min. The TCA precipitates were washed twice with 10 ml acetone and resuspended in 2 ml of 2-N-morpholinoethanesulfonic acid (MES) buffer (pH 5.0). The protein content was measured using the Bradford assay with bovine serum albumin (BSA) as the calibration standard. The total DNA concentration was measured by fluorimetry using the fluorescent dye 4,6-diamidino-2-phenylindole (DAPI) (Jiao et al., 2010) after extraction using Soil DNA purification kit (HiMedia). Comparative profiling of eDNA content was performed by PCR amplification of 16SrRNA gene using primers VC16SF: GTAAAGTACTTTCAGTAGGGAGGA and VC16SR: CATTTGAGTTTTAATCTTGCGACC. The total carbohydrate content was measured using a modified phenol-sulfuric acid method (Jiao et al., 2010). Briefly, 50 µl of EPS solution was mixed with 125 µl of concentrated sulfuric acid. Then 25 µl of 10% phenol was mixed in, and the mixture was incubated in a 95°C water bath for 5 min. The absorbance at 490 nm was recorded. Small glycolipids were quantified by a modified orcinol method. Briefly, 600 ml EPS solution was extracted three times with 600 ml diethyl ether. The diethyl ether extracts were vacuum-concentrated to dryness by evaporation of the organic solvent. The lipids were dissolved in 100 ml deionized water, and 100 ml of 1.6 % (w/v) orcinol and 800 ml 60 % (v/v) H_2_SO_4_ were added. After incubation at 80°C for 30 min and subsequently at room temperature for 10 min, the OD of the solution was measured at 420 nm.

### Antibiotic sensitivity assay, MIC Determination and viability of biofilms

Susceptibility to antimicrobial agents was determined by the disc (Himedia) diffusion method according to the CLSI (The Clinical and Laboratory Standards Institute) guidelines. A number of disc concentrations were used to measure the zones of inhibition after 18 to 24 hrs of incubation at 37°C. Antibiotics indicating interaction with triterpenoids were used for MIC determination using standard protocol. Preformed biofilms exposed to triterpenoids were treated with specific antibiotics at 1X MIC concentration. CFU for the biofilms were enumerated by resuspending the film and plating on antibiotic/ triterpenoid free medium.

### Assaying EPS associated enzyme activities

Phosphatase specific activity was measured by the release of p-nitrophenol (PNP) from p-nitrophenyl phosphate as described previously (Macaskie et al., 2000). Activity was expressed as % of activity compared to untreated samples. Qualitative assay of protease was performed in media plates containing 1% gelatin in a solution of 1.5% agar. Biofilm-culture supernatants filtered with 0.22µm filter were poured into wells bored onto the media and incubated for overnight. Diameters of clear zones were measured to evaluate protease activity of the culture filtrates (Montville, 1983). Proteolytic activity in biofilm-culture supernatants was assayed using azocasein as substrate for quantitative assay. Digestion was done in 20 mM Tris/HCl buffer (pH 8.0) containing 10 mM CaCl_2_. The amount of digested substrate was measured at 440 nm. Activity was expressed as % of activity compared to untreated samples. Exolipase activity was assayed using p-nitrophenylpalmitate as the substrate. 10 ml of isopropanol containing 30 mg of p-nitrophenylpalmitate was mixed with 90 ml of 0.05M Sorensen phosphate buffer (pH 8.0) containing 207 mg of sodium deoxycholate and 100 mg of gum arabic. A 2.4 ml amount of this freshly prepared substrate solution was prewarmed at 37°C and then mixed with 0.1 ml of cell free supernatant fluid. After 15 min of incubation at 37°C, the OD was measured at 410 nm against an enzyme-free control. Activity was expressed as % of activity compared to untreated samples.

### Hydrophobicity of the cell surface

A 1.2 ml volume of cell suspension in phosphate/urea/Mg (PUM) buffer, pH 7.5 (22.2 g K_2_HPO_4_. 3H_2_O, 7.26 g KH_2_PO_4_, 1.8 g urea, 0.2 g MgSO_4_. 6H_2_O, 1 l deionized water), was mixed with 200 ml of n-hexadecane with vigorous stirring for 2 min. After phase separation at room temperature for 15 min, the lower aqueous phase was removed and its OD measured at 580 nm.

### Homology model and Molecular docking

The structure of VcVpsT, VhLuxP-LuxQ and VcHapR were obtained from PDB (PDB ID: 3kln, 1zhh and 2PBX). PyMOL v1.3 was used to visualize the structural models (Peters et al., 2006). *In silico* docking of structural models of VpsT, LuxP, LuxQ and HapR were performed with GRA, UA and BA using Molecular docking simulation PyRX (Dallakyan and Olson, 2015) consisting of AutoDockvina (Trott and Olson, 2010). The binding site residues were identified from the LigPlot+ (Laskowski and Swindells, 2011) for representation of hydrophobic and hydrogen-bond interactions.

## Results

Prior to determining the impact on biofilm formation, MBC for the triterpenoids were determined. Log-phase bacterial cells (OD ~ 0.6) was exposed to the phytochemicals for concentration range of 0 to 1000 μg/ ml. The MBC of the triterpenoids were determined by calculating CFUs against log phase bacterial cultures after 16 h (over-night) of triterpenoid treatment (GRA, UA and BA). The MBC values against *V. cholerae* was determine to be > 1000 μg/ ml, > 400 μg/ ml and > 400 μg/ ml for GRA, UA and BA respectively (Table 1). Sub-MBC concentration ranges of 0-400 μg/ ml for GRA and 0 – 200 μg/ ml for both UA and BA were chosen with an upper limit of 0.5XMBC for subsequent analysis. Bacterial biofilm formation was estimated by conventional crystal violate staining with overnight cultures treated with selected concentrations of triterpenoids (100μg/ ml, 200μg/ ml and 400μg/ ml for GRA and 50μg/ ml, 100μg/ ml and 200μg/ ml for UA and BA). As depicted in Fig. 1, against *V. cholerae*, the three phytochemicals exerted analogous influence on biofilm formation. For GRA a gradual decrease of biofilm formation was observed from 100μg/ ml (69.94%) which further reduced at higher concentrations (33.82% for 200μg/ ml and 25.89% for 400μg/ ml, Fig. 1A). Compared to untreated cells, UA treated cultures exhibited reduced biofilm formation (65.75% for 50μg/ ml, 44.71% for 100μg/ ml, 21.92% for 200μg/ ml, Fig. 1B). Similarly, BA showed considerable effect on the *V. cholerae* biofilm formation as well with increasing concentrations (63.33% for 50μg/ ml, 24.97% for 100μg/ ml, 13.98% for 200μg/ ml, Fig. 1C). The influence of the triterpenoids was further envisioned at higher concentrations. When tested at higher concentrations, GRA and UA demonstrated reduction in biofilm formation around 3-fold for 800 μg/ ml and 400 μg/ ml respectively (P<0.001), however at 400 μg/ ml, BA enhanced biofilm formation by 1.5 fold (P < 0.01) (Fig. S1). Preformed (24h, static) biofilms were assessed for dispersion after treatment with the compounds. Dispersion remained unaffected by GRA and BA treatment however with UA slight enhancement of dispersion was noted (1.6-times compared to control at 200 μg/ ml; P < 0.05) (Figs. 1D, 1E and 1F). Viability of cells in biofilms, formed in presence of the triterpenoids, were also tested where CFU value decreased dose dependently for GRA, UA and BA, reinforcing the antibiofilm activity of the triterpenoids (9.5, 14.06 and 10.9 – fold reduction for 400 μg/ ml of GRA or 200 μg/ ml of UA or BA treatment respectively) (Figs. 1G, 1H and 1I and Fig. S2). Colony morphology, particularly rugose and smooth colony-type transition has been associated with biofilm forming potential. In order to examine if triterpenoids can affect colony morphology, C6709 cells were spotted on plates containing 400 μg/ ml of GRA or 200 μg/ ml of UA or BA and allowed to form mature colony for 72 h. As presented in Fig. 1J, BA and, to a lesser extent, UA significantly reduced rugosity of the colony compared to untreated or GRA treatment. Since swarming motility have been linked with biofilm formation in *V. Cholerae* (Seghal Kiran et al., 2016), we tested the triterpenoids for their possible impact on swarming motility on semisolid (0.5%) agar plates. Interestingly, all three teriterpenoids displayed enhanced swarming motility increasing bacterial swarm diameter by 3.06-fold, 2.58-fold and 1.83-fold respectively with GRA, UA and BA (Fig. S3 and Figs. 2A, 2B and 2C). For determining the impact of the triterpenoids on auto-aggregation, the property often linked with seeding and maturation of biofilms, C6709 cells were treated in presence and absence of 400 μg/ ml of GRA or 200 μg/ ml of UA or BA for overnight; were allowed to form auto-aggregate at room temperature in PBS for 3hrs and subsequently analysed by carefully measuring the OD_600_ for non-aggregated cells. Interestingly, only GRA affected auto-aggregation of the bacteria (12.2%) compared to the untreated control (28.73%). UA and BA on the flip-side apparently did not affect auto-aggregation (Fig. 2D). The result indicates potential of the three triperpenoids to interfere at various stages of biofilm maturation. Ability of the triterpenoids to affect adherence on glass surface was determined after allowing mid-log phase culture of *V. cholerae* to adhere on glass for 3 h. Bacterial adherence was visualized by CV-staining of the surface. As depicted in Fig. 2D, 200 μg/ ml for UA and BA; 400 μg/ ml for GRA severely impaired surface adherence of the cells. Intriguingly, despite their apparent consistency in perturbing bacterial biofilms, the precise impact on dynamics of biofilm formation might be variegated for the three molecules. While the three triterpenoids affected adherence of planktonic cells on glass substratum, it is only GRA that affected auto-aggregation of planktonic C6709 cells. To further assess the impact of the three triterpenoids on the integrity and architecture of the biofilms, Scanning Electron Microscopic (SEM) studies were performed. Biofilm formation was allowed in absence and presence of the three triterpenoids (100 μg/ ml for UA and BA; 200 μg/ ml for GRA) and the resulting biofilm formed after 24hrs of exposure was viewed under SEM. Micrographs showed thick meshed-network of biofilm for untreated cells but GRA, UA and BA clearly demonstrated considerable loss of the integral cellular network (Figs. 3A, 3B, 3C and 3D). The EPS production was visibly impaired in the all the treated samples (Fig.3), however size and shape of individual cells remained apparently unaffected.

**Table 1.** Determining the test concentrations on the basis of MBC value for test-compounds.

| **Triterpenoid** | **MBC vs C6709 (µg/ ml)** | **Maximum test conc. (µg/ ml)** |
| --- | --- | --- |
| **GRA** | >1000 | 400 |
| **UA** | 400-500 | 200 |
| **BA** | 400-500 | 200 |

**Fig. 1.**
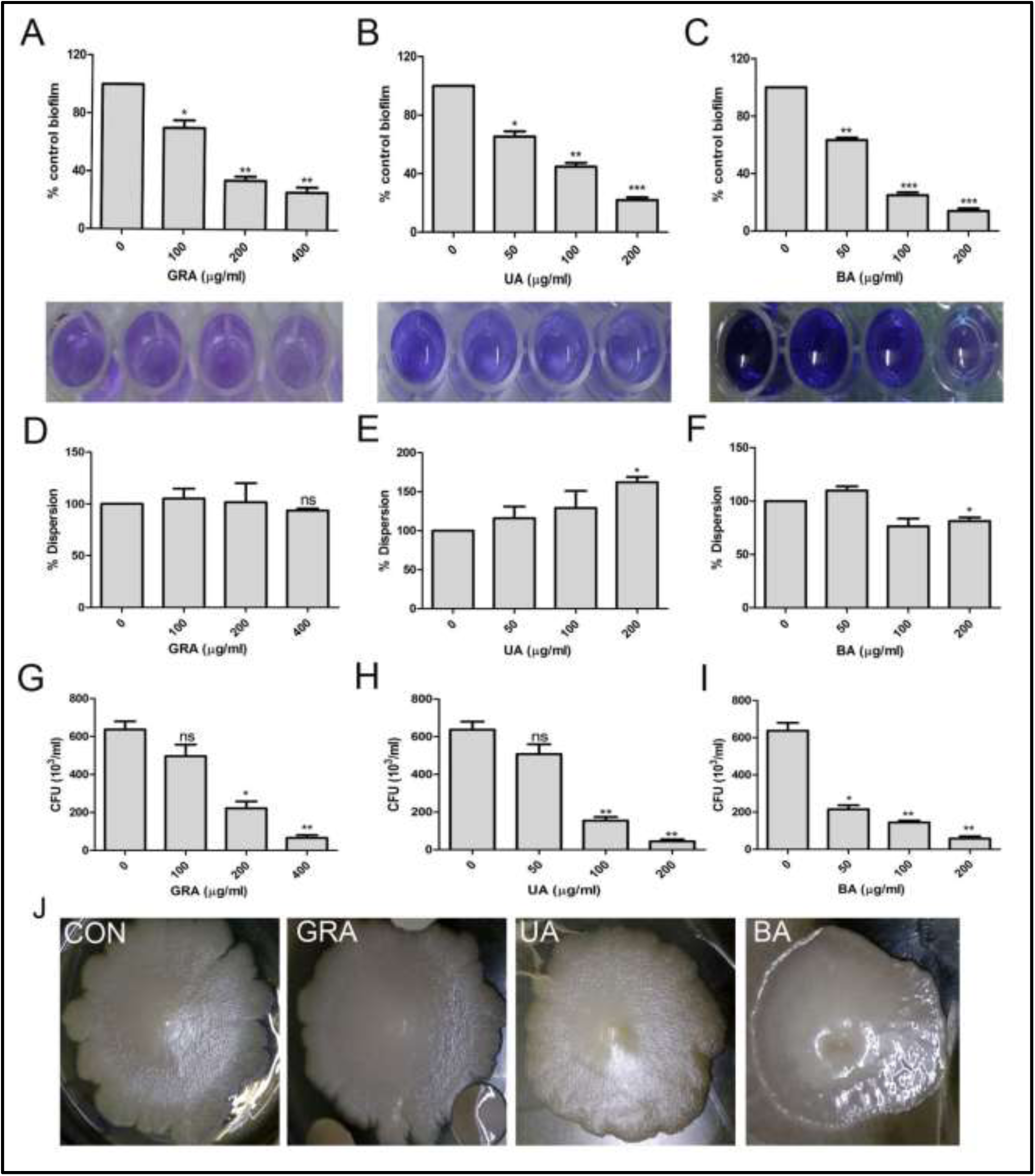
Effect of triterpenoids on biofilm formation and dispersion. Log phase cultures of *V. cholerae* C6709 was allowed to form biofilm in presence of various concentrations of GRA (A), UA (B) and BA (C) and the formation of biofilm was evaluated by crystal violate (CV) staining. Lower panel: stain retention following resuspension in 70 % ethanol. Preformed biofilms were exposed to GRA (D), UA (E) and BA (F) and dispersion of biofilmed cells were examined. Viability of biofilmed cells formed in presence of various concentrations of GRA (G), UA (H) and BA (I) was assessed by determining CFU from each system. (J) Surface texture and colony morphology of C6709 cells grown in absence (CON) or in presence of GRA (400 μg/ ml); UA (200 μg/ ml) and BA (200 μg/ ml) were visualized after 72 h of inoculation. Data are representative of at least three independent replicates. *P < 0.05; **P < 0.01; ***P < 0.001; ns not significant, two tailed paired student t-test.

**Fig.2.**
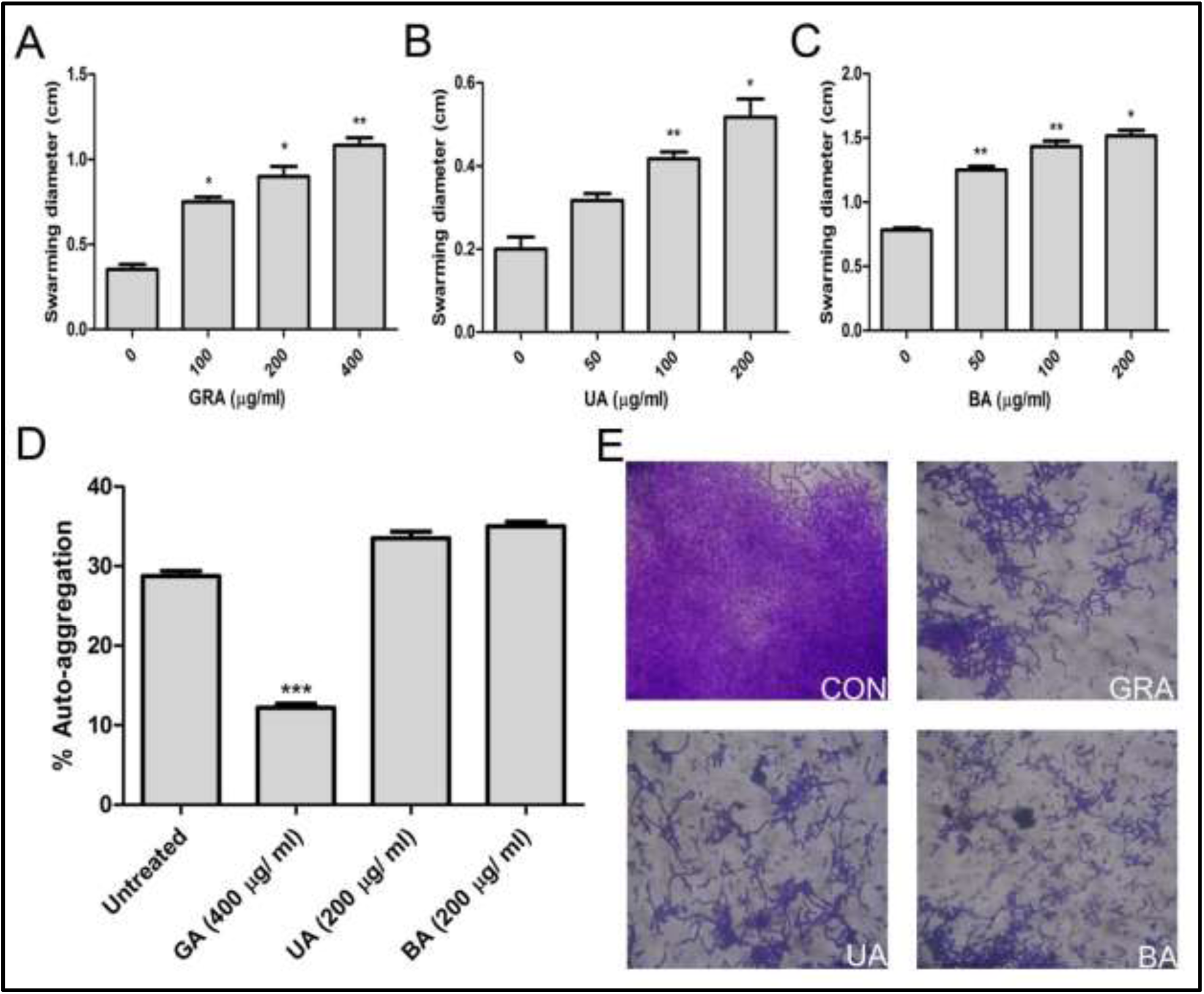
Impact of triterpenoid exposure on motility and adherence. Swarming motility of *V. cholerae* C6709 cells were evaluated by measuring swarming diameters on semisolid medium in presence of various concentrations of GRA (A), UA (B) and BA (C). (D) Late log phase C6709 cells were treated with GRA (400 μg/ ml); UA (200 μg/ ml) and BA (200 μg/ ml). Auto-aggregation was quantified for each treatment. (E) Adherence of C6709 cells on glass substratum was observed by CV-staining after allowing the cells to adhere on glass surface for 3 h in absence (CON) or in presence of GRA (400 μg/ ml); UA (200 μg/ ml) and BA (200 μg/ ml). Data are representative of at least three independent replicates. *P < 0.05; **P < 0.01; ***P < 0.001, two tailed paired student t-test.

**Fig.3.**
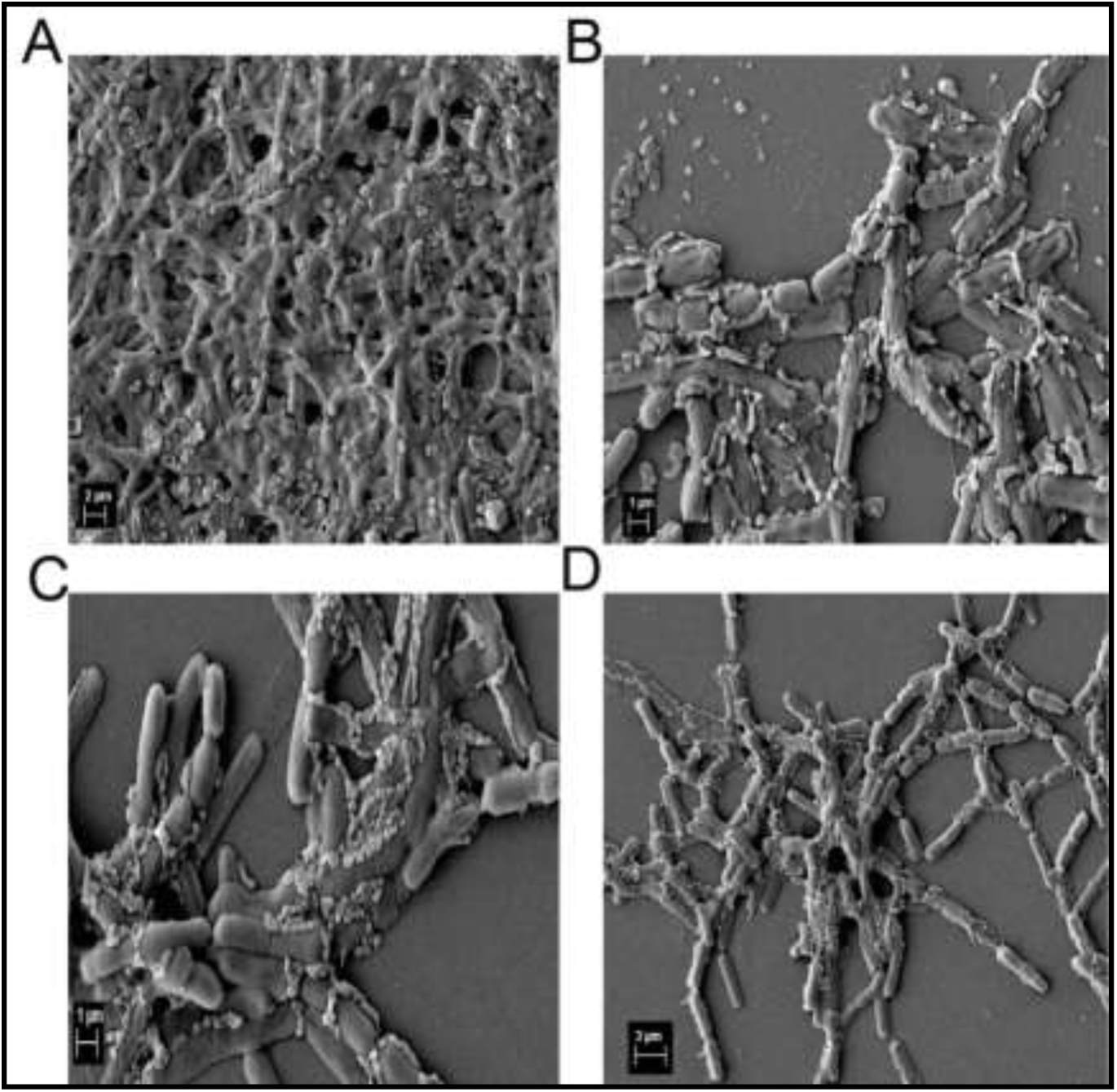
Triterpenoids severely impaired the structural integrity of biofilms. C6709 cells were allowed to form biofilm on glass substratum for 24 h in absence (A) or in presence of GRA (400 μg/ ml) (B); UA (200 μg/ ml) (C) and BA (200 μg/ ml) (D). Scanning electron microscopic observation of C6709 biofilms formed were recorded. Representative images of at least 25 fields from each system are shown.

The biofilm extracellular matrix is composed of *Vibrio* polysaccharide (VPS) consisting primarily of glucose, galactose and proteins to coat substratum surface for adhesion (RbmC and Bap1) and enhancing cell-cell cohesion (RbmA) (Berk et al., 2012; Fong and Yildiz, 2007). Since the BA, UA and GRA affected various aspects of biofilm formation including EPS production as indicated by SEM, comprehensive physico-chemical profiling of C6709 biofilm was performed. For visualizing possible alteration of gross chemical composition, EPS was extracted from biofilms developed in presence or absence of the three triterpenoids (400 μg/ ml of GRA or 200 μg/ ml of UA or BA) and analyzed by FT-IR spectroscopic scan. As represented in Fig.4A, cluster of peaks corresponding to –COC-groups were detectable around 3400 cm^-1^ (preak1) and amide1 from proteins were detectable 1647 cm^-1^. For peak 1 several signals were detectable in untreated control, while the signals diminished with UA and BA treatment. For GRA treatment few of the signals were detected. For peak 2, 1.5%, 4% and 10.1% increase in % transmittance was recorded (Fig. 4A). The results hinted towards alteration of EPS composition by the triterpenoid exposure. For specific physico-chemical analysis, EPS extraction was performed from biofilms grown on semi solid medium in presence or absence of UA (200 μg/ ml), BA (200 μg/ ml) or GRA (400 μg/ ml). GRA, UA and BA reduced carbohydrate content of EPS by 28%, 18.56% and 16.50% respectively compared to control (Fig. 4B). The slime production during biofilm development was studied by growing the biofilms on 30 µg/ ml congo red agar plates. Substantial mitigation of congo red accumulation was observed in biofilms developed on 200 μl/ ml BA containing medium (Fig. S4). Considerable reduction in total protein content of EPS was recorded with GRA, UA and BA resulting in 1.88-fold, 1.97-fold and 3.59-fold reduction in total precipitable proteins (Fig. 4C). Glycolipid biosurfactants produced by specific *Vibrio* strains anatagonistically affect biofilms from other strains. Total glycolipid content was measured with chloroform-methanol extracted fraction of EPS by orcinol which showed ~2-fold reduction for each of the treatments compared to untreated controls, primarily indicating depletion of glycolipid content (Fig. 4D). The amount of extracellular DNA, which orchestrates EPS integrity (Okshevsky and Meyer, 2015) was estimated by measuring DAPI-fluorescence (Fig. 4E) and by amplifying *V. cholerae* 16S rRNA gene from eDNA samples prepared after biofilm formation post treatment with GRA (400μg/ ml), UA 200μg/ ml) and BA (200μg/ ml) (Fig. 4F). Significant depletion of eDNA was observed in C6709 biofilms formed in presence of the triterpenoids with BA maximally reducing eDNA content in the EPS (Figs. 4E and 4F). Cell-surface hydrophobicity is a major physical attribute, often associated affinity toward substratum (Mizan et al., 2016). Hydrophobicity of the C6709 cell surface were significantly reduced (1.6-1.8-fold) when grown in presence of GRA, UA or BA (Fig. 5A).

**Fig.4.**
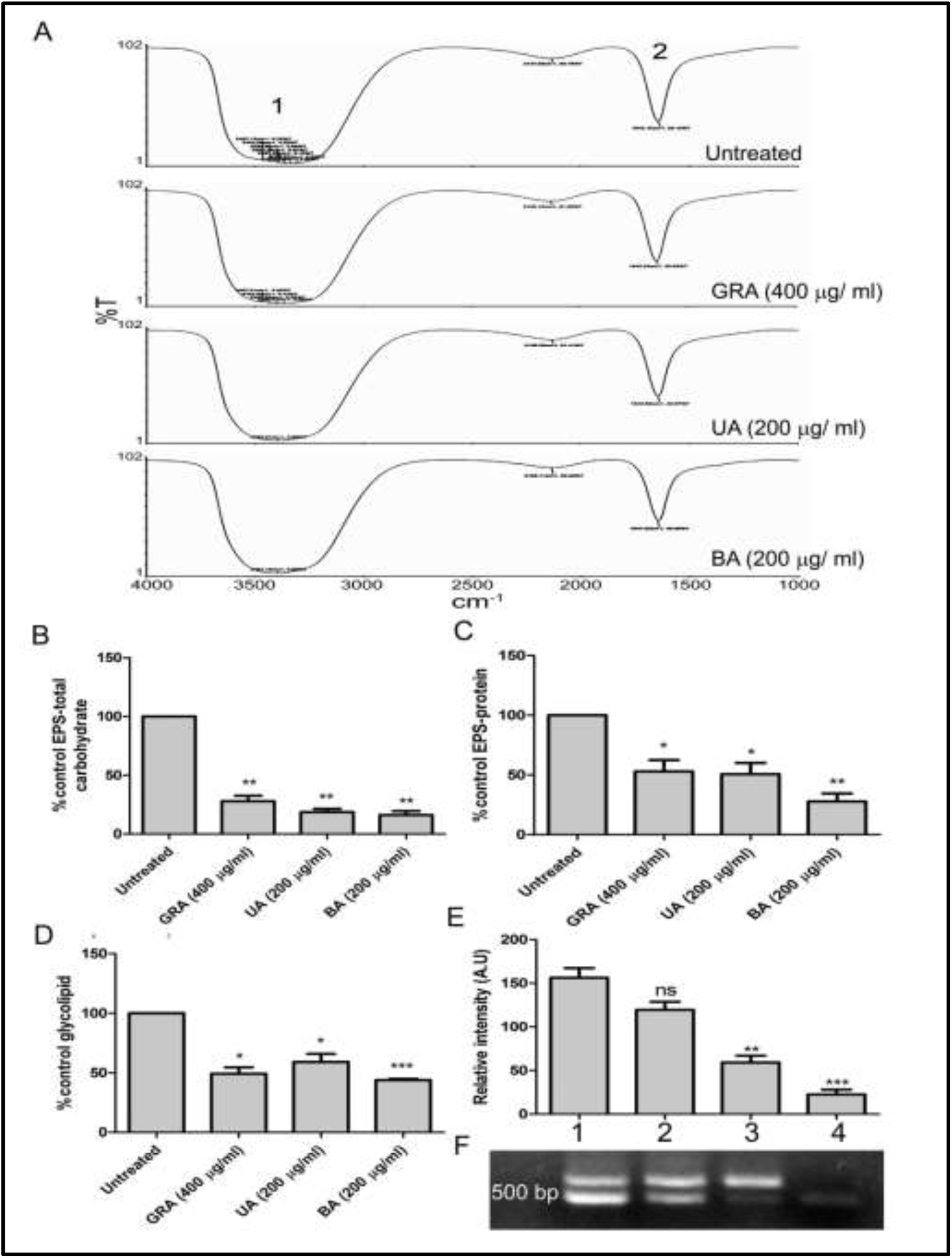
Alteration of EPS composition by triterpenoid treatment. (A) FTIR spectra of EPS from biofilms developed in absence and in presence of GRA (400 μg/ ml); UA (200 μg/ ml) and BA (200 μg/ ml). Peak 1 (cluster), —COC— group vibrations in carbohydrates, DNA, and RNA, peak 2, amide I in proteins. Comparative profile of total carbohydrate (B), protein (C) and glycolipid (D) content quantified as described in materials and methods. (E) eDNA was quantified detecting DAPI fluorescence signal from EPS preparation from untreated (1), GRA (400 μg/ ml) (2); UA (200 μg/ ml) (3) and BA (200 μg/ ml) (4) and by semi quantitative PCR targeting 16S RNA genes (F). Data representative of at least three independent replicates. *P < 0.05; **P < 0.01; ***P < 0.001; ns not significant, two tailed paired student t-test.

**Fig.5.**
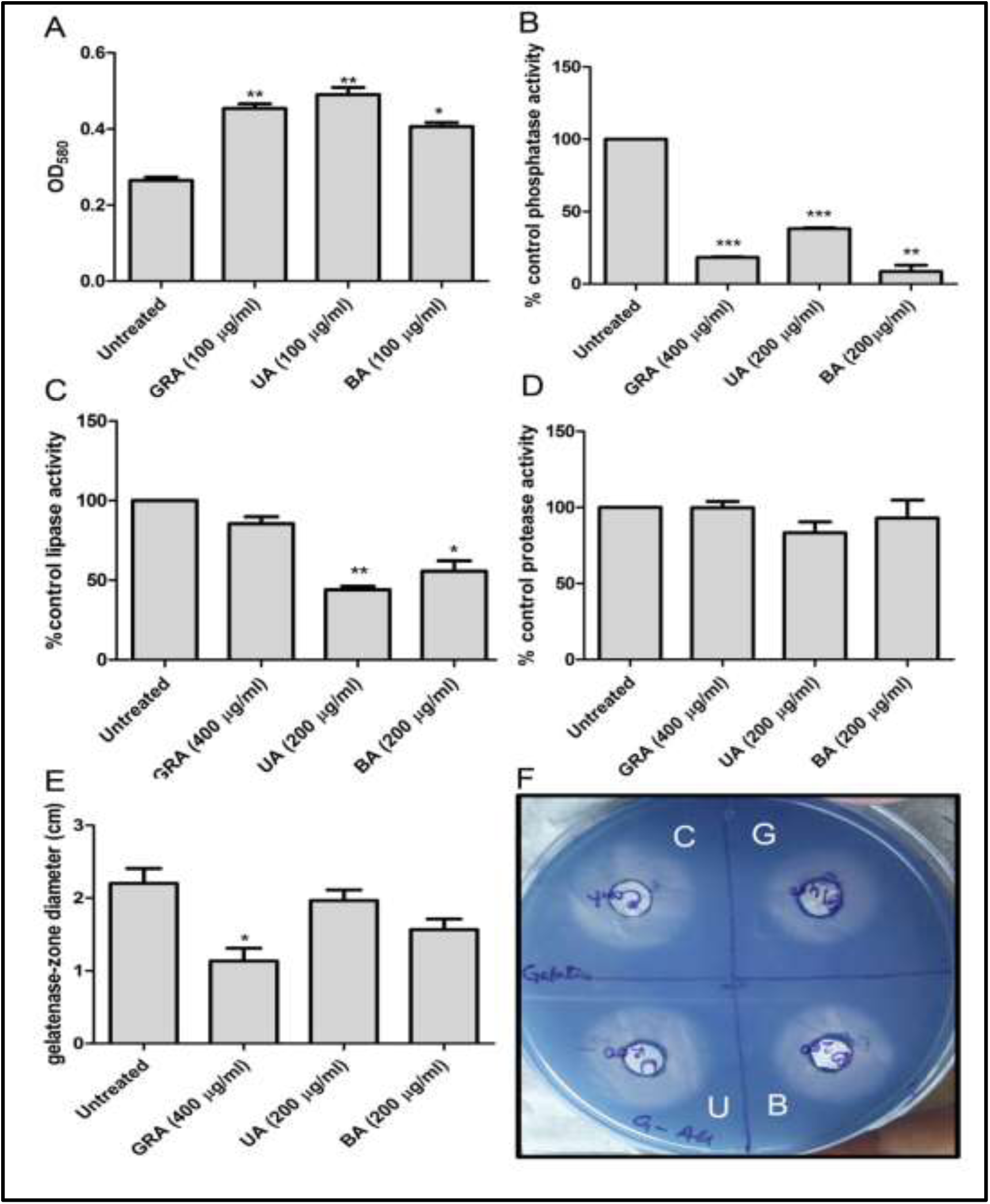
Cell surface hydrophobicity and extracellular enzyme activity. (A) Cell surface hydrophobicity was measured with log-phase C6709 cells treated with GRA (100 μg/ ml); UA (100 μg/ ml) and BA (100 ug / ml). (B) Total extracellular phosphatase activity was determined by pNpp hydrolysis in EPS prepared from C6709 biofilms developed in presence of GRA (400 μg/ ml); UA (200 μg/ ml) and BA (200 μg/ ml) alongside untreated control. (C) Extracellular lipase activity determined by para-nitrophenylpalmitate hydrolysis. (D) Total EPS associated protease activity was determined by azocasein assay and gelatinase activity estimated by proteolytic zone diameter on gelatin agar plates (E). (F) Representative coomassie stained gelatinase assay plate depicting biofilm associated activity in untreated (C), GRA (400 μg/ ml) (G); UA (200 μg/ ml) (U) and BA (200 μg/ ml) (B). Data are representative of at least three independent replicates. *P < 0.05; **P < 0.01; ***P < 0.001, two tailed paired student t-test.

To attain a deeper insight into the impact of the triterpenoids on biofilm physiology, EPS-associated enzyme activity was profiled. GRA, UA or BA were found to substantially diminish the phosphatase activity in the EPS preparations (Fig. 5B). Extracellular lipase activity was also assayed using para-nitrophenylpalmitate as substrate. UA (100µg/ ml) resulted in significant reduction of lipase activity (2.2-fold). BA (100µg/ ml) also resulted in statistically significant decrease in lipase activity (Fig. 5C). The triterpenoids seemed not to affect EPS associated protease activity as reflected by azocasein hydrolysis based assay (Fig. 5D). However when the assay was performed on gelatin agar plate, GRA was found to reduce gelatinase activity by 2-fold (P<0.05). UA and BA did not exhibit significant impact on gelatinase activity of the EPS preparations (Fig. 5E).

Since antibiofilm agents are often associated with enhancement of antibiotic action against bacterial biofilms, interaction between the triterpenoids and series of antibiotics were studied. For preliminary screening of antibiotic-triterpenoid interaction, conventional Kirby-Bauer method for determining antibiotic responsiveness was adopted with discs adsorbed with various antibiotics on plates with or without triterpenoids. While BA was found to augment action of two antibiotics from fluoroquinolone family, namely ofloxacin and ciprofloxacin, GRA and UA significantly enhanced inhibitory potential of representatives from cephalosporin group, namely cefotaxime, cefepime and ceftriaxone (Tables S2 and S3 and Fig. 6A).To test whether presence of the triterpenoid can indeed affect antimicrobial potential against planktonic cells, MIC for the antibiotics and antibiotic-triterpenoid combinations (ofloxacin and ofloxacin with BA; Ciprofloxacin and ciprofloxacin with BA and cefepime, cefotaxim and ceftriaxone with or without UA or GRA independently) were determined by both macrodilution and microdilution. Considerable impact on MIC was determined for UA as cefepime, cephtriaxone and cefotaxim showed reduction of 2, 2.2 and 1.8-fold respectively. GRA displayed lesser impact on the MIC of the antibiotics (Figs. 6B and 6C). As depicted in Fig. 6D, the MIC for ciprofloxacin and ofloxacin was reduced by 1.5 fold when exposed in presence of BA. To explore if such interactions are indeed translated in the action of the antibiotics against biofilmed C6709 cells, cefotaxim was selected for possible interaction with GRA (400 µg/ ml) and UA (200 µg/ ml) while ciprofloxacin was studied for possible interaction with BA (200 µg/ ml). Preformed biofilms were exposed to the antibiotics at various concentrations in combination with respective triterpenoids and CFU for each of the exposed combinations were determined. UA appeared to enhance antibiotic mediated killing effectively at all the tested cefotaxim concentrations with 4.75, 5.6 and 9.0-fold reduction in CFU values at 2.5, 5 and 10 µg/ ml respectively. For GRA-cefotaxim combination significant impact was detected at 10 µg/ ml with 6.7-fold reduction in CFU count (Figs. 6E and 6F). BA-ciprofloxacin combination showed 3-4-fold reduction in CFU counts at all the tested concentrations of ciprofloxacin (0.5, 1.0 and 2.0 µg/ ml) (Fig. 6G). With an objective to validate candidate antibiotic-triterpenoid combinations, we profiled the impact of GRA, UA and BA on responsiveness of a series of antibiotics against planktonic cells. Although the approach could only suggest us potential antibiotic-triterpenoid combination, the augmentation of antibiotic action was further quantified by determining MIC of antibiotic-triterpenoid combination. GRA and UA could potentiate action of cephalosporin class of β-lactam antibiotic. It is not clear whether GRA and UA can induce uptake of cephalosporin group of antibiotics or interacts with penicillin binding proteins to enhance cephalosporin action. On the contrary, BA potentiated two fluoroquinolones, namely ciprofloxacin and ofloxacin. Fluroquinolones target topoisomerases and BA is a known inhibitor of eukaryotic topoisomerase I (Chowdhury et al., 2002), further exploration of fluroquinolone-BA combination might shed light on bacterial topoisomerase inhibition.

**Fig. 6.**
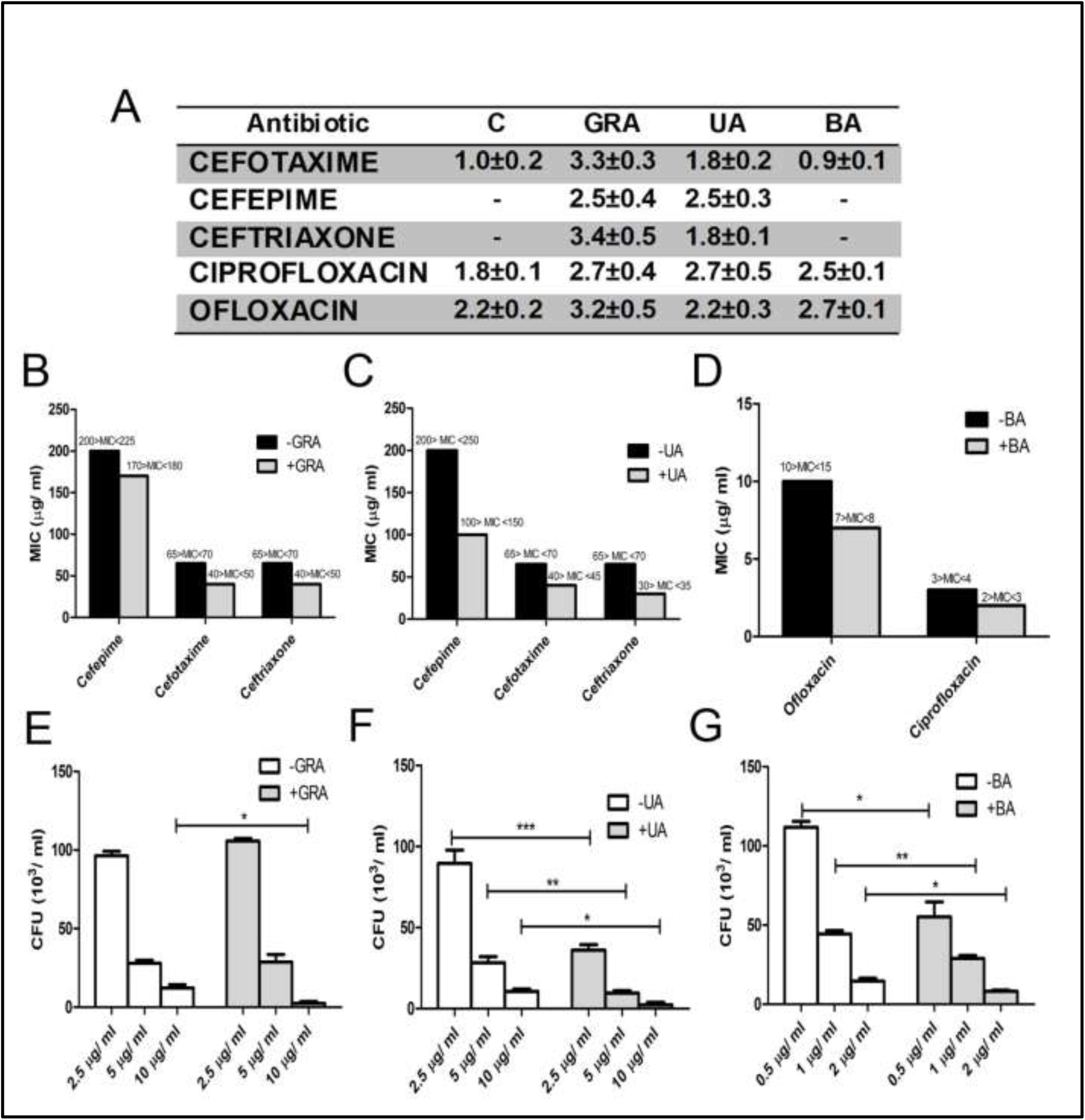
Impact of triterpenoids on antibiotic action. (A) Summary of Kirby–Bauer assay (Table S2 and S3) for detecting impact of triterpenoids on antibiotic action. Combinations suggesting potentiation of antibiotic action are listed. (B, C) Minimum inhibitory concentrations (MIC) of cefepime, cefotaxim and ceftriaxone against C6709 cells were determined in absence or in presence of GRA (400 μg/ ml)(B) or UA (200 μg/ ml) (C). (D) MICs for ofloxacin and ciprofloxacin against C6709 cells as determined without or with BA (200 μg/ ml). (E, F and G) Preformed biofilms were treated with various concentrations of antibiotic and triterpenoids. Viability of biofilmed cells were enumerated by determining CFU of biofilms exposed to cefotaxim-GRA (400 μg/ ml) (E); cefotaxim-UA (200 μg/ ml) (F) and ciprofloxacin-BA (200 µg/ ml) (G) combinations. Data are representative of at least three independent replicates. *P < 0.05; **P < 0.01; ***P < 0.001, two tailed unpaired student t-test.

Intervention of biofilm formation, in particular for Gram negative bacteria, has often been associated with development of molecules that perturb N-acylatedhomoserine (ASL) triggered quorum sensing (QS) mechanism (Ball et al., 2017). Since QS on the contrary to other Gram negative pathogens, negatively regulates biofilm formation in *Vibrio* where co-regulation of multiple QS systems culminates into modulation of biofilm forming and virulence gene expression (Liu et al., 2017). In concert, the QS systems regulate two transcription factors, HapR and AphA, to control the genes for biofilm and virulence regulation in an opposite manner. Crystal structure of LuxP-LuxQ and HapR has been determined from *V. harveyi* and *V. cholerae*. To assess the possibility of GRA, UA and BA to affect QS systems of *Vibrio*, molecular docking was performed with LuxP, LuxQ and HapR. GRA, UA and BA were docked with the dimeric interface of HapR with ΔG of −8.8, −8.1 and −9.1 kcal/ mol (Figs. S5A, S5B and S5C). The interactions identified were primarily hydrophobic interaction with three polar contacts (Figs. S5D, S5E and S5F) suggesting possible impact on the transcriptional regulator. LuxP and LuxQ are one of the earlier identified QS-receptor sensor kinase system triggered by AI-2. GRA, UA and BA docked with LuxP and LuxQ within a hydrophobic groove of the proteins primarily with hydrophobic interactions with 2-3 polar contacts with ΔG around −8 kcal/ mol (Figs. S6 and S7). The cyclic di-GMP sensor VpsT regulates biofilm matrix formation (Krasteva et al., 2010). GRA, UA and BA were docked with monomeric VpsT with ΔG of −7.4, −7.0 and −7.1 kcal/ mol (Figs. 7 A, 7B and 7C). The interactions involved are predominantly hydrophobic with 2-3 polar contacts. These results indicate possibilities of the triterpenoids to function as a modulator of QS in *V. cholerae*. Independent experiments in our laboratory using *Chromobacterium violacium* and *P. aeruginosa* revealed that each of the three pentacyclic triterpenoid can inhibit ASL dependent QS (Paul *et al*., unpublished). However, further efforts are needed to pinpoint the mechanism of triterpenoid mediated modulation of QS.

**Fig. 7.**
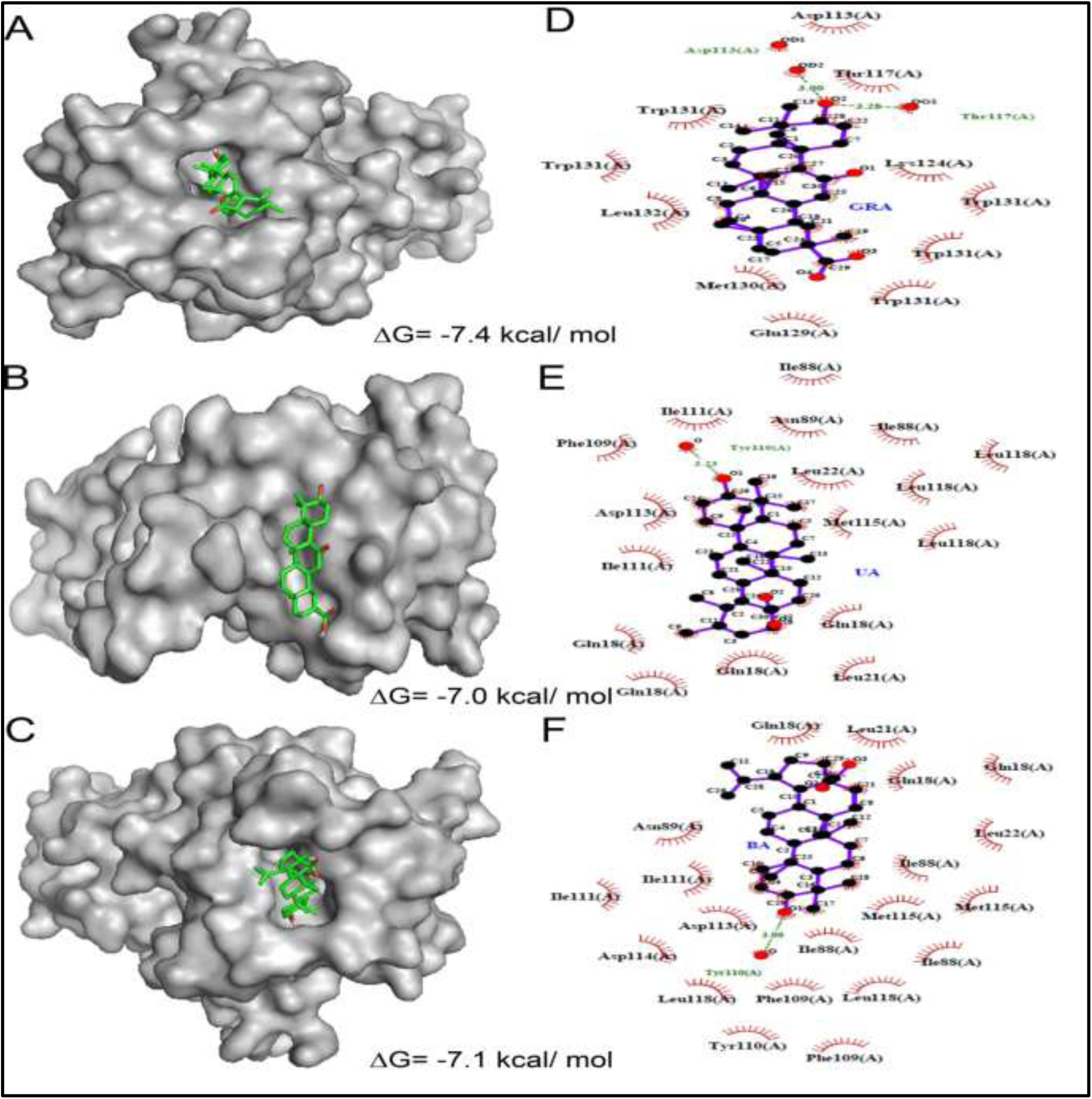
Molecular docking predicts possible interaction of the triterpenoids with the VpsT. The monomeric structure of VpsT was availed from PDB (3KLN) and molecular docking was performed with GRA, UA and BA. The models depicting interaction of VpsT with GRA (A), UA (B) and BA (C). The amino acid residues are represented as a surface with green (Carbon), blue (Nitrogen) and red (Oxygen). Atoms from GRA, BA or UA are represented in ball and stick with purple (Carbon), blue (Nitrogen), red (Oxygen). The binding site residues were identified from the LigPlot+ representations illustrating hydrophobic and hydrogen-bond interactions with GRA (D), UA (E) and BA (F).

## Conclusion

The aim of our work was to comprehensively evaluate potential application of penatcyclic triterpenoids as auxiliary antimicrobial agent. In this study, we considered *V. cholerae* C6709 cells as a model biofilm former and tested commercially available purified form of three triterpenoids representing distinct groups of the phytochemical. Though biofilm formation in *V. cholerae* was primarily associated with transmission, presence of both planktonic cells and biofilm-like structures in exudates from cholera patients underlined the significance of biofilm intervention during infection (Faruque et al., 2006) . In the last decade, many reports have demonstrated the potent antibacterial activities of triterpenes including particularly UA, GRA and BA (Horiuchi et al., 2007). However, majority of the works often analysed molecules purified from specific plant sources and hence there are apparent discrepancy in antimicrobial efficacy of each of the compounds. In corroboration with the study conducted by Fontanay et al., (2008) (Fontanay et al., 2008), our analysis could not detect substantial antibacterial activity for the molecules with MBC values > 500 µg/ ml. Moreover, one major issue regarding clinical implication of the observations can be the potential cytotoxic impact of triterpenoids on various mammalian cell lines. However, the triterpenoids tested in this study can provide the bioactive scaffold to enhance biofilm selectivity and neutralizing cytotoxicity. Moreover on-going efforts to developing nano-delivery strategy for biofilms can be an effective alternative (Mehrizi et al., 2018) to enhance the therapeutic potential for the molecules. Confirming synergy with antibiotic and developing an optimal triterpenoid-antibiotic composition can lead to development of combinatorial therapeutics exploiting these bioactive molecules. Over all, comprehensive understanding of anti-biofilm potential of pentacyclic triterpenoids can offer the scope of developing multi-faceted strategies to combat bacterial infections.

## Acknowledgments

The authors acknowledge Prof. Sanjay Ghosh, Dept. of Biochemistry, University of Calcutta for his guidance and providing laboratory facilities when needed. The authors are thankful to Prof. Tapas Sengupta and Scanning Electron Microscopy division of IISER-Kolkata for their support. The authors also acknowledge Dr. Rukhsana Chowdhury, CSIR-IICB, Kolkata for kindly providing C6709 strain and offering crucial suggestions. The authors would like to specially mention the FTIR facility of IACS, Kolkata for their constant support.

## Author contributions

AS, SPB and AB designed the experiments. SPB and AB performed the experiment. AS, SPB and AB analysed the data and prepared the manuscript. All authorised and approved the final draft.

## Compliance with ethical standards

