## Supplemental file 1 for "A comprehensive and comparative study on the action of pentacyclic triterpenoids on *Vibrio cholerae* biofilms"

**Supportive file:**

Fig. S1. **Effect of triterpenoids at elevated dose on biofilm formation**

Fig. S2. **Viable cell count from biofilms formed with triterpenoids**

Fig. S3. **Titerpenoids enhanced swarming motility**

Fig. S4. **BA significantly reduced slime formation from C6709 biofilms.**

Fig. S5. **Molecular docking predicts possible interaction of the triterpenoids with HapR**

Fig. S6. **Molecular docking predicts possible interaction of the triterpenoids with the LuxQ**

Fig. S7. **Molecular docking predicts possible interaction of the triterpenoids with the LuxQ**

Table S1: **Determination of MBC for the triterpenoids.**

Table S2: **Summary of Kirby-Bauer assay on plates containing GRA and UA**.

Table S3: **Summary of Kirby-Bauer assay on plates containing BA**

Table S4: **Data for FTIR scan of EPS preparations (separate MS-Excel file)**

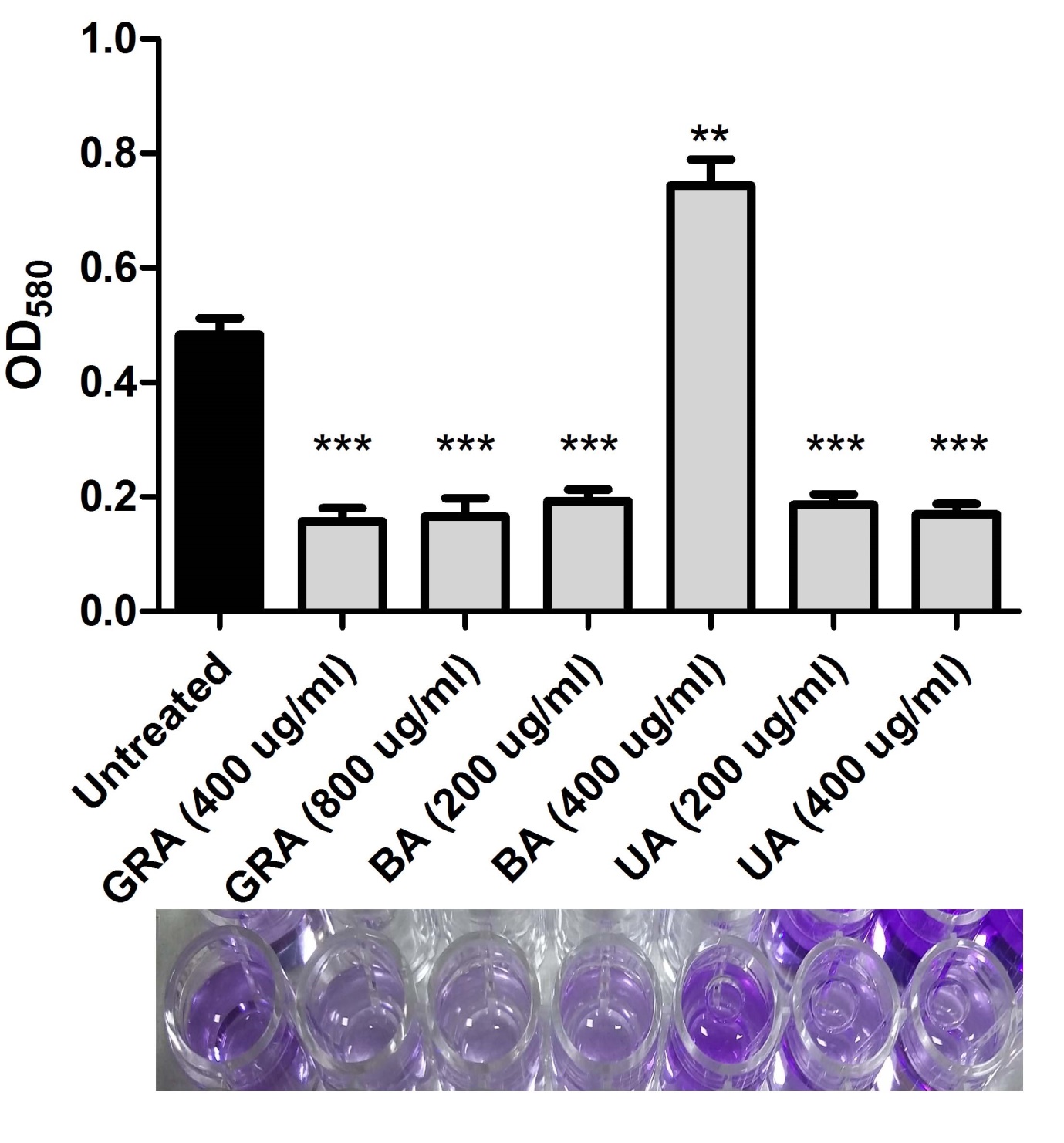

**Fig. S1.** **Effect of triterpenoids at elevated dose on biofilm formation.** Logphase cultures of *V. cholerae* C6709 was allowed to form biofilm in presence of 0.5X and 1X MBC of GRA, UA and BA and the formation of biofilm was evaluated by crystal violate staining. Lower panel: stain retention following resuspension in ethanol. *P < 0.05; **P < 0.01; ***P < 0.001, two tailed unpaired student t-test.

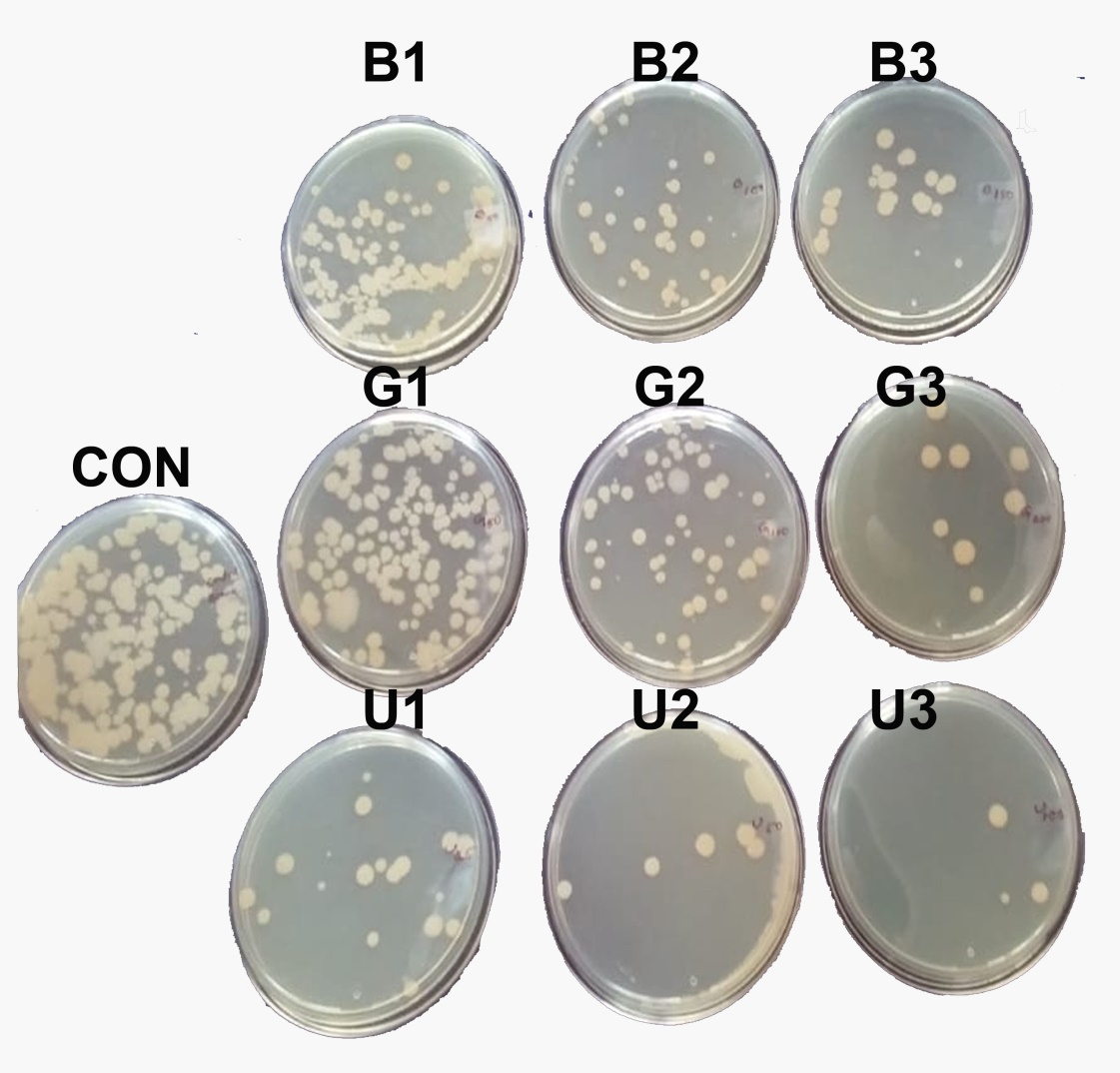

**Fig. S2.** **Viable cell count from biofilms formed with triterpenoids.** Biofilms derived from GRA (G1, G2 and G3 for 100, 200 and 400 μg/ ml), UA (U1, U2 and U3 for 50, 100 and 200 μg/ ml) or BA (B1, B2 and B3 for 50, 100 and 200 μg/ ml) exposed systems were enumerated for viable cell count (CFU) and compared with CFU for untreated control (CON).

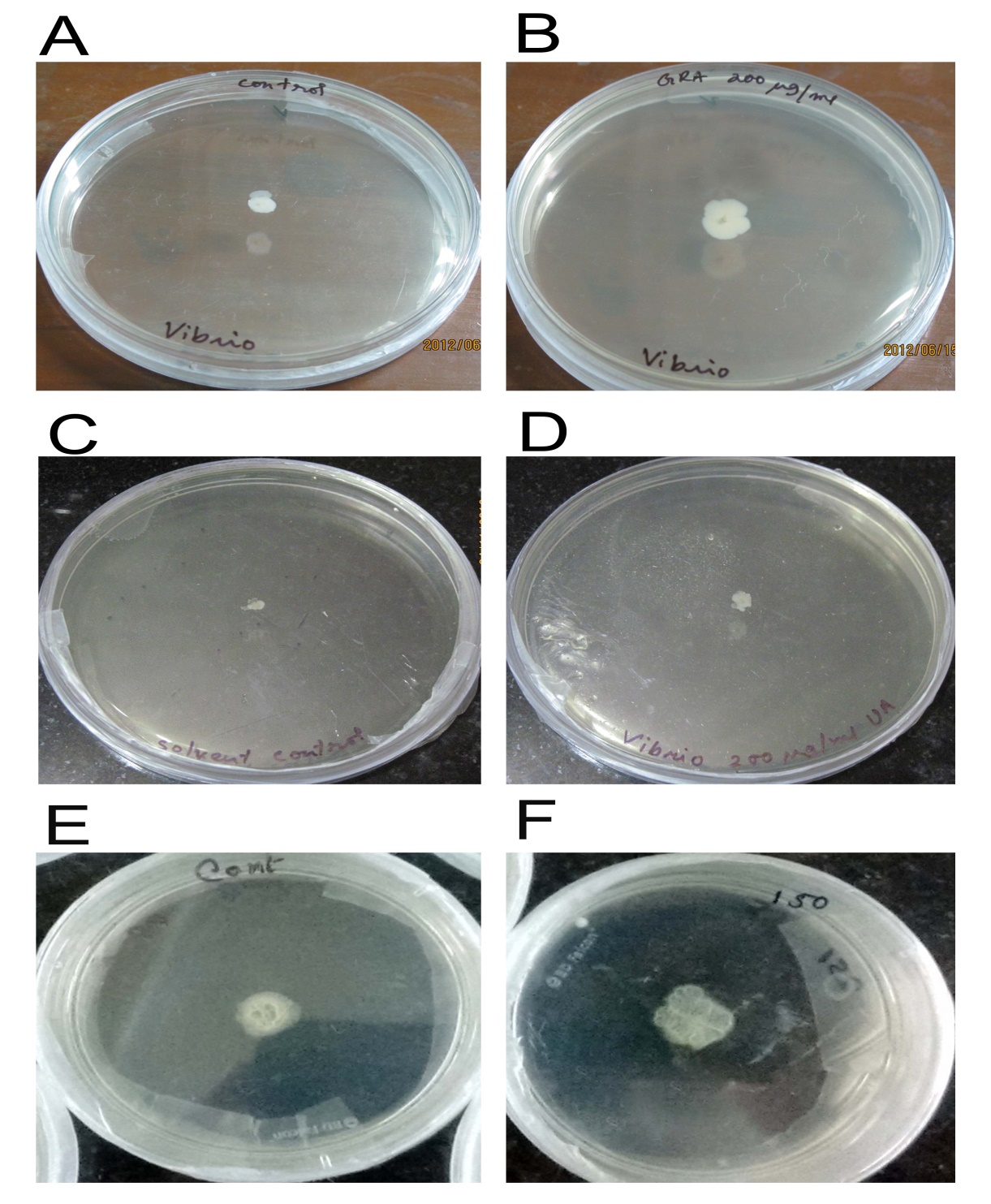

**Fig. S3**. **Titerpenoids enhanced swarming motility.** Log phase C6709 cells were spotted on semisolid plates with out or with GRA (400 μg/ ml); UA (200 μg/ ml) or BA (200 μg/ ml) (A,B; C;D and E,F respectively) and swarming was observed after 36h.

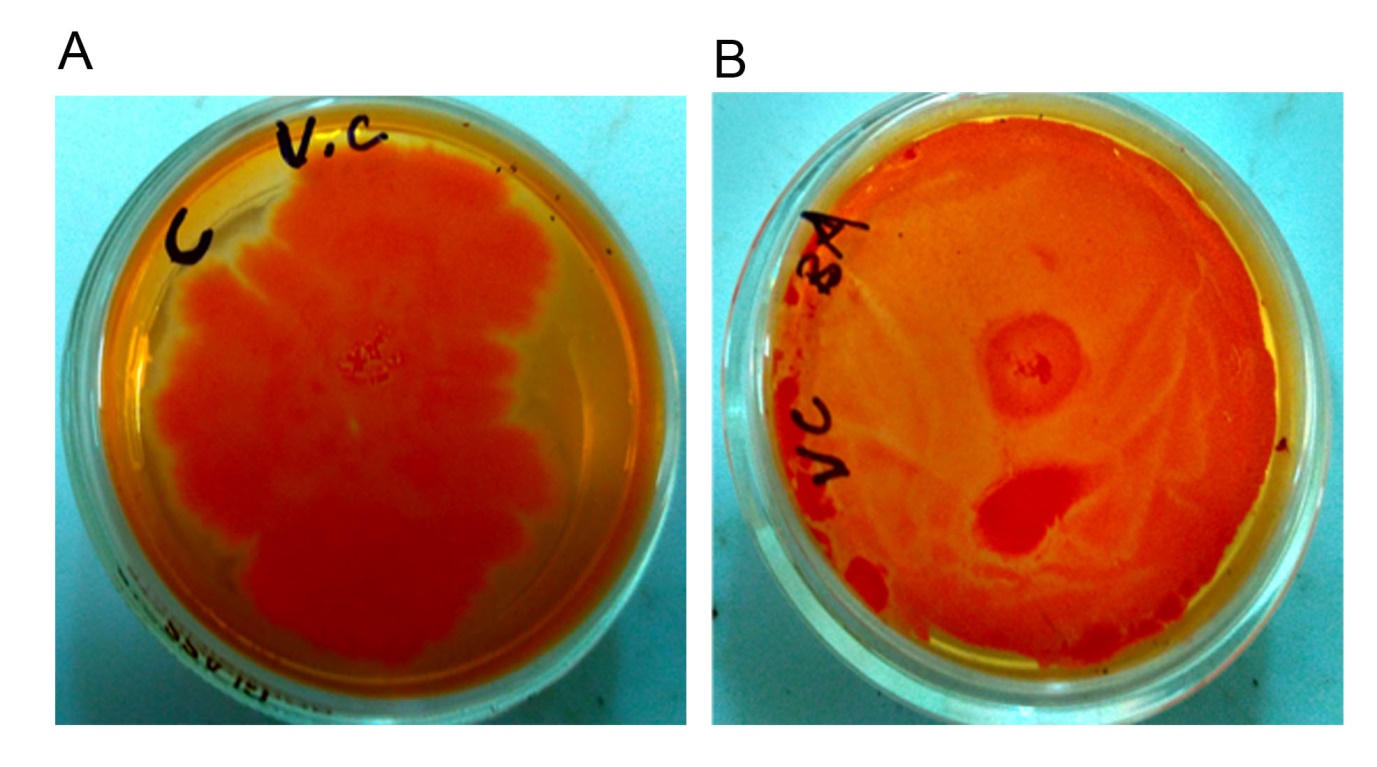

**Fig. S4.** **BA significantly reduced slime formation from C6709 biofilms.** C6709 cells were allowed to form biofilms on congo red agar plates in absence (A) and presence of BA (200 μg/ ml) (B). Trapping of congo red indicates slime formation by individual biofilms.

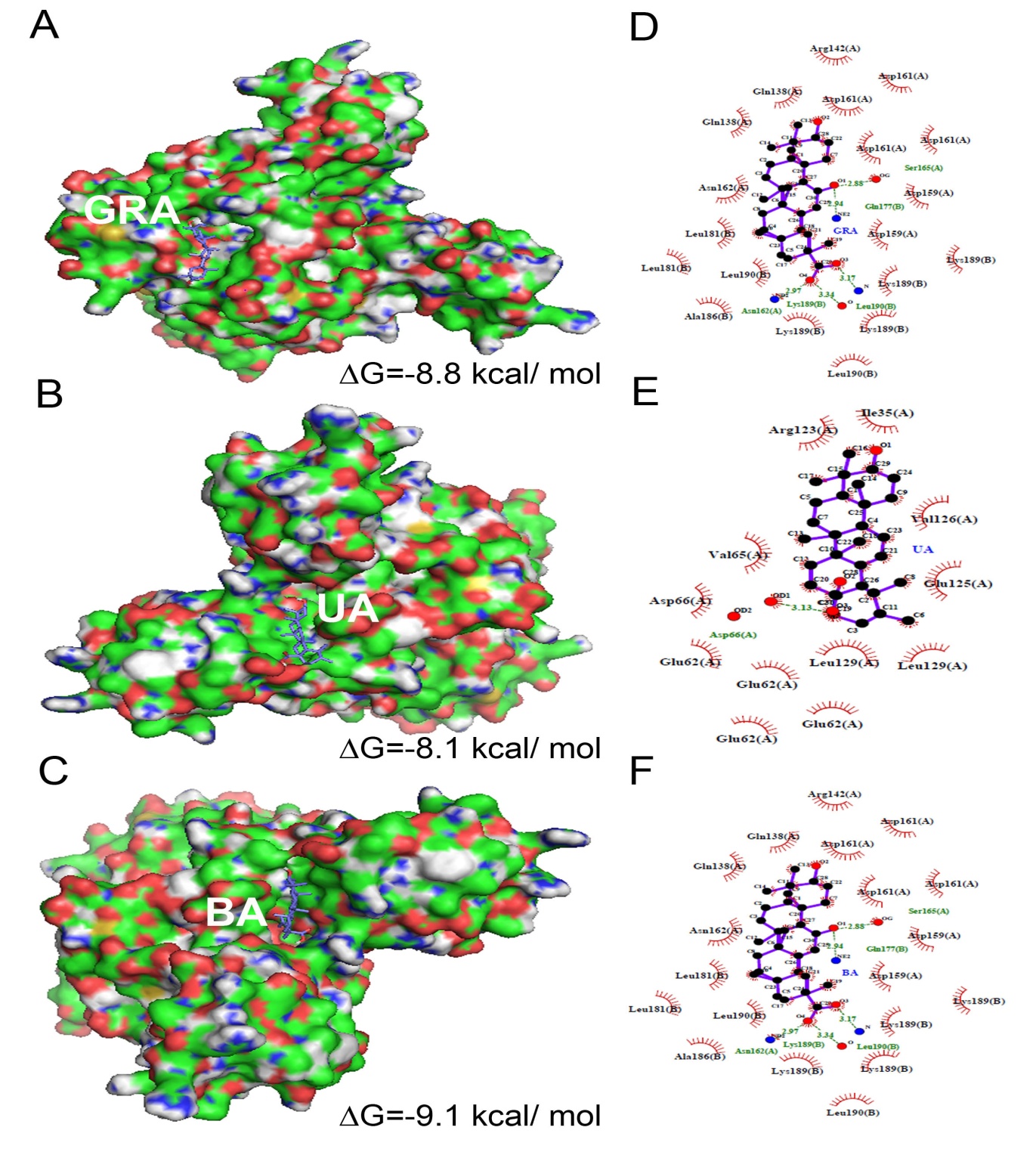

**Fig. S5** Molecular docking predicts possible interaction of the triterpenoids with the HapR. The dimeric structure of hapR was availed from PDB (2PBX) and molecular docking was performed with GRA, UA and BA. The models depicting interaction of HapR with GRA (A), UA (B) and BA (C). The amino acid residues are represented as a surface with green (Carbon), blue (Nitrogen) and red (Oxygen). Atoms from GRA, BA or UA are represented in ball and stick with purple (Carbon), blue (Nitrogen), red (Oxygen). The binding site residues were identified from the LigPlot+ representations illustrating hydrophobic and hydrogen-bond interactions with GRA (D), UA (E) and BA (F).

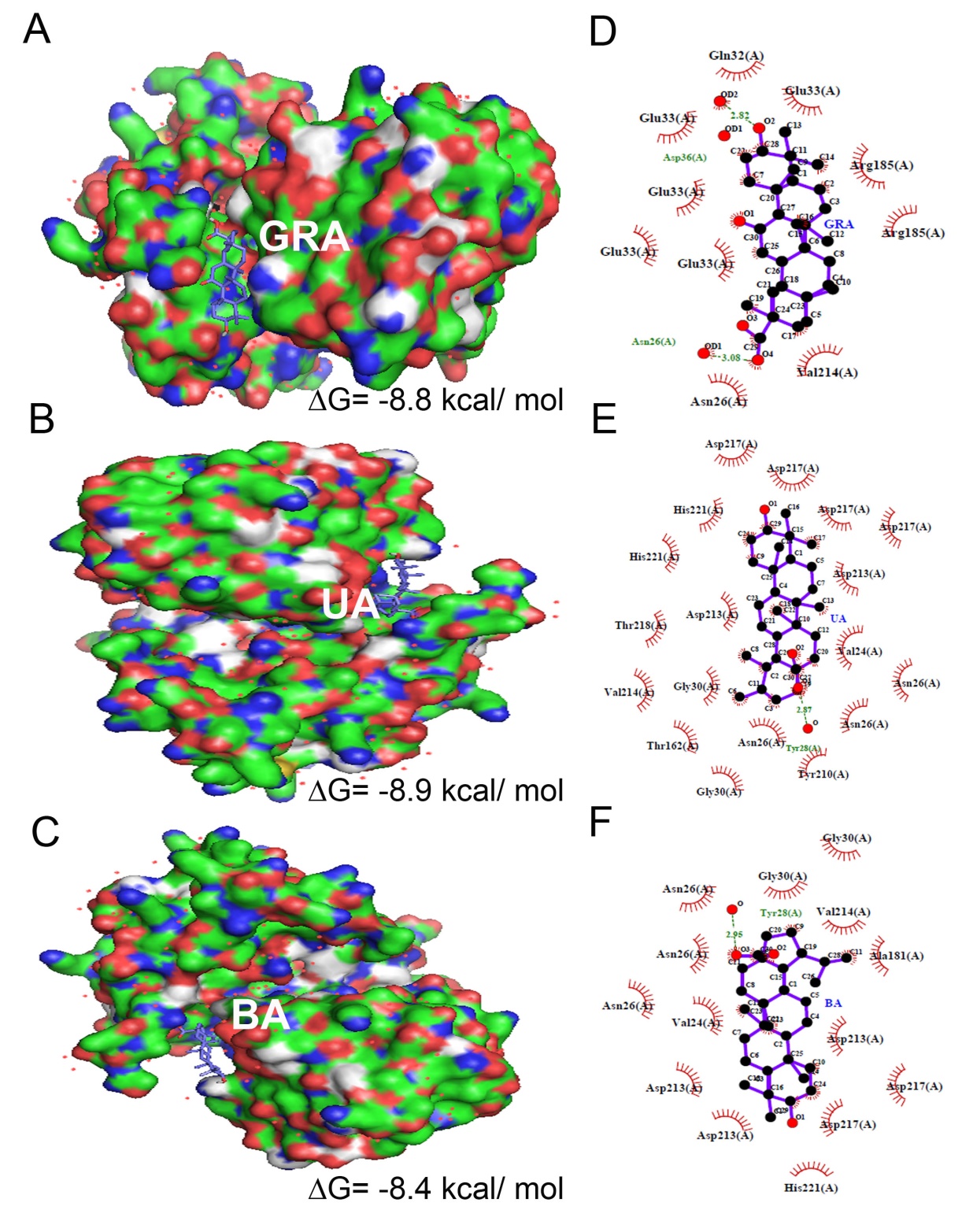

**Fig. S6.** **Molecular docking predicts possible interaction of the triterpenoids with LuxP.** The dimeric structure of LuxP from *V. harveyi* was availed from PDB (1ZHH, chain A) and molecular docking was performed with GRA, UA and BA. The models depicting interactioin of HapR with GRA (A), UA (B) and BA (C). The amino acid residues are represented as a surface with green (Carbon), blue (Nitrogen) and red (Oxygen). Atoms from GRA, BA or UA are represented in ball and stick with purple (Carbon), blue(Nitrogen), red (Oxygen). The binding site residues were identified from the LigPlot+ representations illustrating hydrophobic and hydrogen-bond interactions with GRA (D), UA (E) and BA (F).

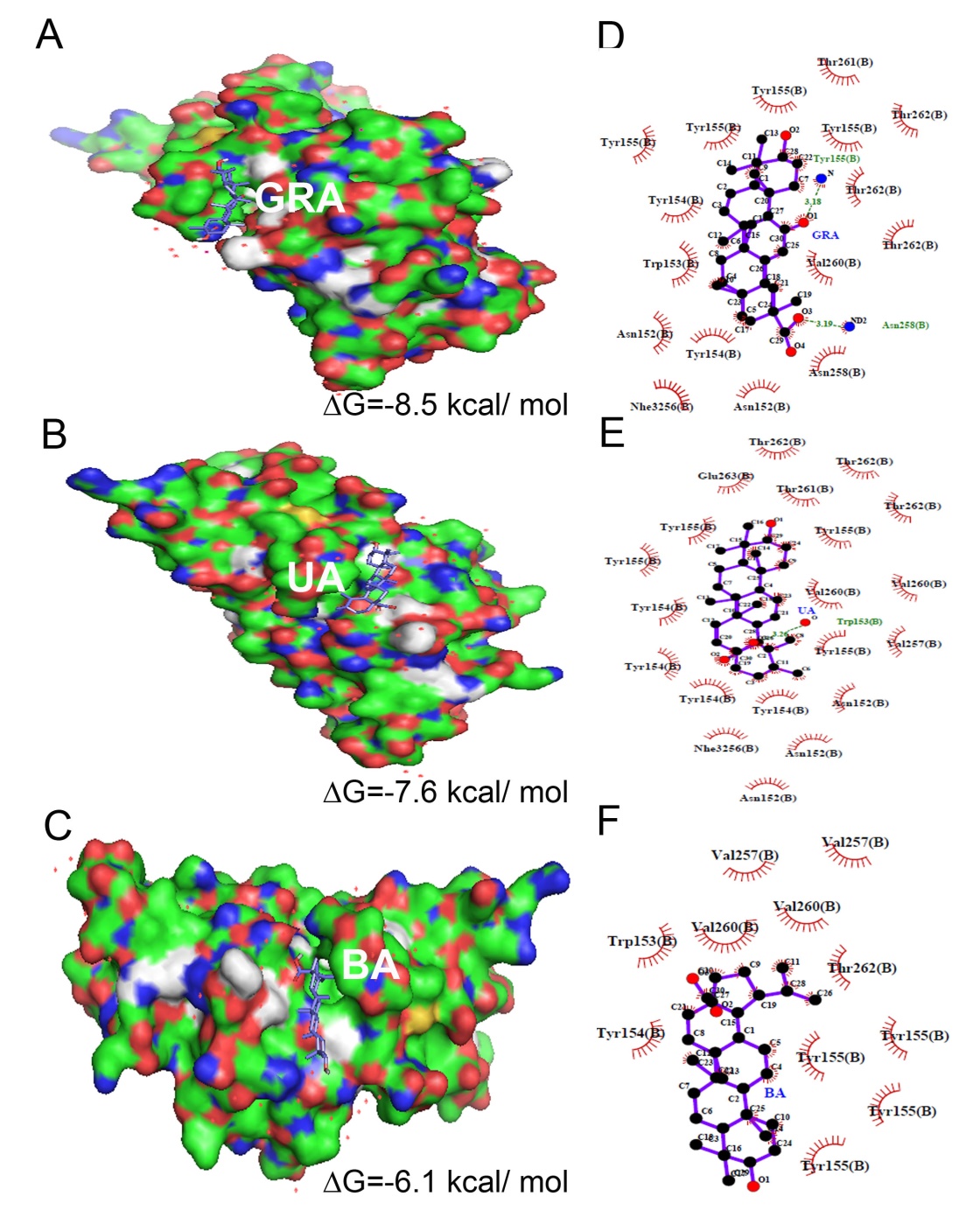

**Fig. S7.** **Molecular docking predicts possible interaction of the triterpenoids with the LuxQ.** The dimeric structure of LuxQ from *V. harveyi* was availed from PDB (1ZHH, chain B) and molecular docking was performed with GRA, UA and BA. The models depicting interaction of HapR with GRA (A), UA (B) and BA (C). The amino acid residues are represented as a surface with green (Carbon), blue (Nitrogen) and red (Oxygen). Atoms from GRA, BA or UA are represented in ball and stick with purple (Carbon), blue(Nitrogen), red (Oxygen). The binding site residues were identified from the LigPlot+ representations illustrating hydrophobic and hydrogen-bond interactions with GRA (D), UA (E) and BA (F).

**Table S1: Determination of MBC for the triterpenoids.** MBCs against *V. cholerae* C6709 by CFU counts following GRA, UA and BA exposure. Observations from three independent experiments are recorded.

| Conc. (μg/ ml) | UA | BA | GRA |
| --- | --- | --- | --- |
| 0 | lawn | lawn | lawn |
| 50 | lawn | lawn | lawn |
| 100 | lawn | lawn | lawn |
| 150 | 200-300 | 200-300 | 200-300 |
| 200 | 200-300 | 200-300 | 200-300 |
| 250 | 100-200 | 100-200 | 200-300 |
| 300 | 100-200 | 100-200 | 200-300 |
| 350 | 50-100 | 50-100 | 100-200 |
| 400 | 50-100 | 50-100 | 100-200 |
| 450 | 25-50 | 25-50 | 100-200 |
| 500 | <10 | <10 | 50-100 |
| 1000 | 0 | 0 | <50 |

**Table S2:** **Summary of Kirby-Bauer assay on plates containing GRA and UA**. Mean diameter of zones of inhibition for each system is depicted.

| Sl. No. | ANTIBIOTICS |  | Diameter of zones of inhibition (cm) | |
| --- | --- | --- | --- | --- |
|  |  | Control | GRA (400 ug/ ml) | UA (200 ug/ ml) |
| 1 | AMIKACIN | 1.8 | 3 | 1.9 |
| 2 | AMOXYCILLIN | 0.9 | 1.1 | 0.9 |
| 3 | AZITHROMYCIN | 1.9 | 2 | 1.3 |
| 4 | CEFALEXIN | - | - | - |
| 5 | CEFEPIME | - | 2.5 | 2.5 |
| 6 | CEFOTAXIME | 1.6 | 3.3 | 1.8 |
| 7 | CEFTRIAXONE | 1.5 | 3.4 | 1.8 |
| 8 | CIPROFLOXACIN | 2.2 | 3.2 | 2.2 |
| 9 | CLINDAMYCIN | 1.9 | 1.3 | 1.8 |
| 10 | DOXYCYCLINE | 2 | 2.9 | 2 |
| 11 | ERYTHROMYCIN | 1.9 | 1.2 | 1.2 |
| 12 | FRAMYCETIN | 1.4 | 2.3 | 2.2 |
| 13 | GATIFLOXACIN | 2.7 | 3 | 2.2 |
| 14 | GENTAMYCIN | 2.4 | 2.9 | 2.6 |
| 15 | OFLOXACIN | 2 | 2.7 | 2.7 |
| 16 | PIPEMIDIC ACID | 1.8 | 3.5 | 2.3 |
| 17 | ROXYTHROMYCIN | 2.9 | 1.9 | 1.3 |
| 18 | TOBRAMYCIN | 2 | 3.4 | 2.2 |
| 19 | VANCOMYCIN | 1.6 | 1.0 | 1.3 |

**Table S3:** **Summary of Kirby-Bauer assay on plates containing BA**. Mean diameter of zones of inhibition for each system is depicted.

| Sl.no. | ANTIBIOTIC | Diameter of zones of inhibition (cm) | |
| --- | --- | --- | --- |
|  |  | Control | BA (200 μg/ ml) |
| 1 | Amikacin | 2.23 | 2.1 |
| 2 | Amoxycillin | - | - |
| 3 | Azithromycin | 1.8 | 2 |
| 4 | Cefalexin | - | - |
| 5 | Cefepime | - | - |
| 6 | Cefotaxime | 0.93 | 1.03 |
| 7 | Ceftriaxone | - | - |
| 8 | Ciprofloxacin | 1.83 | 2.2 |
| 9 | Clindamycin | 2.3 | 2.33 |
| 10 | Doxycycline | 2.1 | 1.9 |
| 11 | Erythromycin | 1.6 | 1.43 |
| 12 | Framycetin | 1.2 | 1.33 |
| 13 | Gatifloxacin | 2.4 | 2.46 |
| 14 | Gentamicin | 2.2 | 2.23 |
| 15 | Ofloxacin | 2.2 | 2.7 |
| 16 | Pipemidic Acid | 1.9 | 1.9 |
| 17 | Roxithromycin | 2.1 | 2.13 |
| 18 | Tobramycin | 2.1 | 2.2 |
| 19 | Vancomycin | 1.2 | 1.06 |
